# MechRNA: prediction of lncRNA mechanisms from RNA-RNA and RNA-protein interactions

**DOI:** 10.1101/285965

**Authors:** Alexander R. Gawronski, Michael Uhl, Yajia Zhang, Yen-Yi Lin, Yashar S. Niknafs, Varune R. Ramnarine, Rohit Malik, Felix Feng, Arul M. Chinnaiyan, Colin C. Collins, S. Cenk Sahinalp, Rolf Backofen

## Abstract

**Motivation:** Long non-coding RNAs (lncRNAs) are defined as transcripts longer than 200 nucleotides that do not get translated into proteins. Often these transcripts are processed (spliced, capped, polyadenylated) and some are known to have important biological functions. However, most lncRNAs have unknown or poorly understood functions. Nevertheless, because of their potential role in cancer, lncRNAs are receiving a lot of attention, and the need for computational tools to predict their possible mechanisms of action is more than ever. Fundamentally, most of the known lncRNA mechanisms involve RNA-RNA and/or RNA-protein interactions. Through accurate predictions of each kind of interaction and integration of these predictions, it is possible to elucidate potential mechanisms for a given lncRNA.

**Approach:** Here we introduce MechRNA, a pipeline for corroborating RNA-RNA interaction prediction and protein binding prediction for identifying possible lncRNA mechanisms involving specific targets or on a transcriptome-wide scale. The first stage uses a version of IntaRNA2 with added functionality for efficient prediction of RNA-RNA interactions with very long input sequences, allowing for large-scale analysis of lncRNA interactions with little or no loss of optimality. The second stage integrates protein binding information pre-computed by GraphProt, for both the lncRNA and the target. The final stage involves inferring the most likely mechanism for each lncRNA/target pair. This is achieved by generating candidate mechanisms from the predicted interactions, the relative locations of these interactions and correlation data, followed by selection of the most likely mechanistic explanation using a combined p-value.

**Results:** We applied MechRNA on a number of recently identified cancer-related lncRNAs (PCAT1, PCAT29, ARLnc1) and also on two well-studied lncRNAs (PCA3 and 7SL). This led to the identification of hundreds of high confidence potential targets for each lncRNA and corresponding mechanisms. These predictions include the known competitive mechanism of 7SL with HuR for binding on the tumor suppressor TP53, as well as mechanisms expanding what is known about PCAT1 and ARLn1 and their targets BRCA2 and AR, respectively. For PCAT1-BRCA2, the mechanism involves competitive binding with HuR, which we confirmed using HuR immunoprecipitation assays.

**Availability:** MechRNA is available for download at https://bitbucket.org/compbio/mechrna

**Supplementary information:** Supplementary data are available at *Bioinformatics* online.

## 1 Introduction

With the advance of large-scale transcriptome analysis it has become evident that the majority of the human genome is transcribed into RNA (Djebali *et al*., 2012). Out of all currently annotated genes, only a minority is known to code for proteins, while most are believed to be non-coding RNAs (ncRNAs). Beside several small ncRNAs, including small nucleolar RNAs (snoRNAs) and microRNAs (miRNAs), manifold analyses showed that especially long non-coding RNAs (lncRNAs), a designation given to any ncRNA longer than 200 nucleotides, play an important role in cell regulation (Marchese *et al*., 2017). The major classes of lncRNAs include natural antisense transcripts (NATs), promoter-associated ncRNAs (pncRNAs), pseudogenes and long intergenic non-coding RNAs (lincRNAs). They have a variety of known functions influencing transcription, splicing, mRNA stability and translation (Kung *et al*., 2013).

For some lncRNAs the specific mechanism of action is known, however often only isolated examples exist. For many others the precise mechanism still needs to be determined. At the most fundamental level, every lncRNA mechanism involves RNA-RNA interaction and/or RNA-protein interaction (and via proteins, DNA interactions). So in order to model lncRNA mechanisms computationally, algorithms for predicting these kinds of interactions are essential. There are a number of tools to predict RNA-RNA interactions. These follow four general approaches, in order of complexity: hybridization-only (RNAHybrid (Rehmsmeier *et al*., 2004), RNADuplex (Lorenz *et al*., 2011)), sequence concatenation (PairFold (Andronescu *et al*., 2005), RNAcofold (Bernhart *et al*., 2006)), accessibility-based (RNAup (Muckstein *et al*., 2006), IntaRNA2 (Mann *et al*., 2017)) and full joint structure prediction -leading to the first joint free energy model for interacting RNA strands (Alkan *et al*., 2006) and follow up work (piRNA (Chitsaz *et al*., 2009), inRNAs (Salari *et al*., 2010), RIP (Huang *et al*., 2009)). Hybridization-only methods, where only intermolecular base-paring is considered, and sequence concatenation methods, where standard algorithms for secondary structure prediction are applied to the concatenation of the input RNA, are very fast but produce unrealistic interactions. Accessibility-based tools compute the partition function of each input sequence and determine the energy required for any given region to be unpaired. These energies are then used as penalties when predicting hybridizations. At the expense of a little higher complexity, the modelled interactions are much more realistic. Accessibility-based tools are efficient enough to have been successfully applied to prokaryotic sRNA and eukaryotic miRNA target prediction on a transcriptome-wide scale. However, due to the complexity of these algorithms, the problem of predicting lncRNA interactions on a transcriptome-wide scale quickly becomes intractable for any method more complex than hybridization-only predictions.

It is possible to use RNA-RNA interaction prediction software for transcriptome-wide, lncRNA-RNA interaction prediction, through the use of existing tools such as IntaRNA (on a supercomputer (Terai *et al*., 2016)) or by new pipelines such as RISearch2 (Alkan *et al*., 2017). All these approaches need to apply the following steps (not necessarily in order): (1) determine accessible regions on every target sequence (e.g. using Raccess (Kiryu *et al*., 2011) and remove repeat regions); (2) determine “seeds” with perfect complementary and extend each seed with flanking sequences of fixed length; and (3) predict (and refine) the interaction between the lncRNA and each of these sequences (e.g. using IntaRNA or RactIP (Kato *et al*., 2010)). Unfortunately the targets of non-coding RNAs identified through the above approach are typically not very specific. For short non-coding RNAs such as sRNAs and miRNAs, it is possible to improve specificity via sequence conservation (Wright *et al*., 2013, 2014) across species. However this does not extend to lncRNAs, which are typically poorly conserved (Iyer *et al*., 2015). As we will discuss below, one way to improve specificity may be to incorporate RNA-protein interactions with RNA-RNA interactions with RNA-protein interactions.

RNA-protein interactions can be determined experimentally using CLIP-Seq, which is currently the standard protocol for the transcriptome-wide identification of RNA-binding protein (RBP) binding sites. Several protocol variants exist, most notably PAR-CLIP (photoactivatable-ribonucleoside-enhanced CLIP) (Hafner *et al*., 2010) and iCLIP (individual-nucleotide CLIP) (Konig *et al*., 2010). Lately, eCLIP (enhanced CLIP) (Van Nostrand *et al*., 2016) and irCLIP (infrared-CLIP) (Zarnegar *et al*., 2016) have been introduced to further improve protocol efficiency with varying approaches, as discussed in (Uhl *et al*., 2017).

A drawback of CLIP-Seq protocols to identify RBP binding sites is that they naturally rely on the expression of the target transcripts, which is often cell-or tissue-specific, especially in the case of lncRNAs (Brunner *et al*., 2012; Liu *et al*., 2016). Computational prediction of missing binding sites is therefore in high demand. While initial prediction methods such as MEME (Bailey and Elkan, 1994) have relied solely on sequence information, more recent tools like MEMERIS (Hiller *et al*., 2006), RNAcontext (Kazan *et al*., 2010) and GraphProt (Maticzka *et al*., 2014) also incorporate structural information to further improve their predictions.

To our knowledge, no tool exists that integrates both RNA-RNA and RNA-protein interactions. This is crucial for lncRNA interaction prediction since their long length increases the probability of protein binding. The type of RBP, whether it binds to the lncRNA or the target and the location of the RBP relative to the RNA-RNA interaction site can allow inference of the potential lncRNA mechanism.

To solve this problem we propose MechRNA, a pipeline for combining interaction predictions and biological data to discover potential mechanisms. Specifically, this pipeline aims to discover potential mechanisms of an input lncRNA by (1) predicting lncRNA-target interactions using IntaRNA2 with a new feature improving transcriptome-wide performance, (2) identifying RBP binding sites predicted by GraphProt on both the targets and the lncRNA, (3) finding correlation between the lncRNA and targets using TCGA expression or user-provided data, (4) combining this evidence to generate candidate mechanisms, and finally (5) computing joint p-values to select the candidate mechanisms that best explain the observed data.

## 2 Methods

MechRNA has four inputs (Ensembl IDs of lncRNA sequence, target sequences and RBPs, and a list of mechansims) and two modes (screening and hypothesis-driven modes). In screening mode, the user only specifies the lncRNA, and the entire transcriptome with all available RBP models is used to predict all possible mechanisms. Since nothing is known about the relationships between the lncRNA, targets and RBPs, correlation data is used to reduce the number of candidates. Hypothesis-driven mode allows the user to specify any *a priori* information they may have on the lncRNA. For example, a common case would be that the lncRNA was experimentally shown to downregulate a set of targets. In this case, the user would specify a list of all downregulatory mechanisms from those that are available, and the list of suspected targets. From these inputs, MechRNA predicts lncRNA-target interactions, RBP binding sites and determines the most likely mechanism given these interactions. Here we will describe each stage in detail. An overview of the pipeline is shown in Figure 1.

**Fig. 1.**
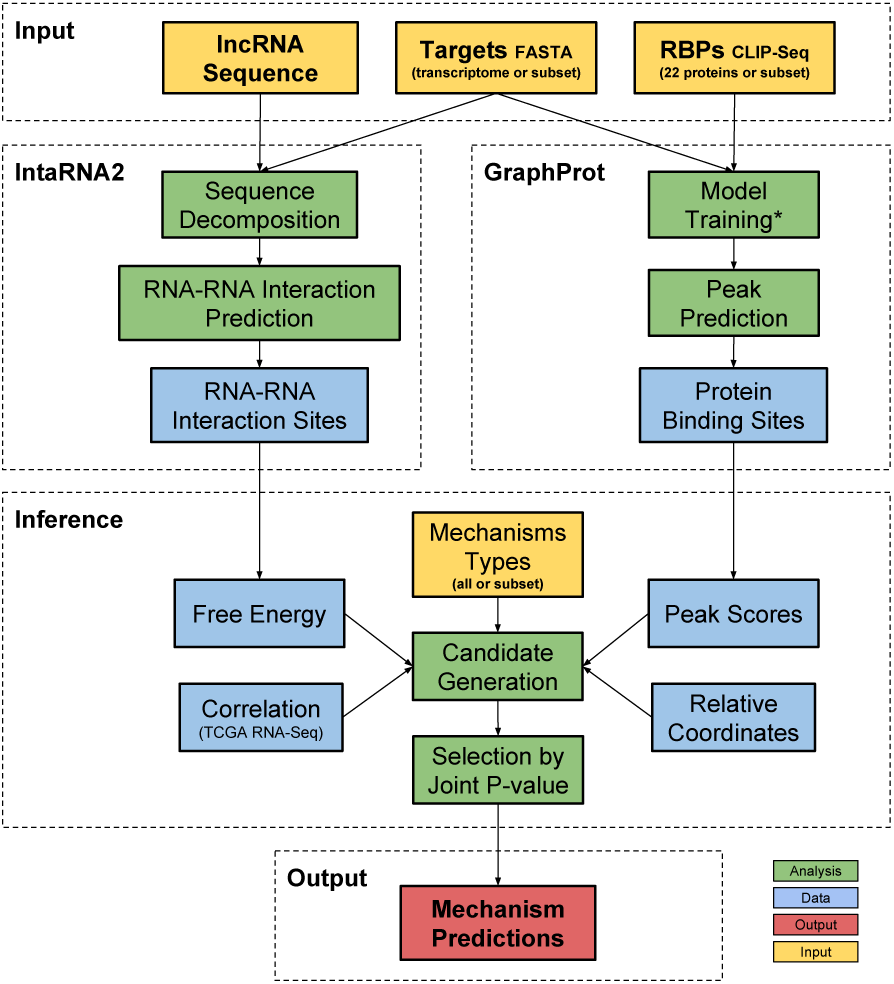
Overview of the MechRNA pipeline. IntaRNA2 computes the optimal RNA-RNA interaction sites between the lncRNA and the accessible regions of targets/transcriptome. GraphProt predicts protein binding sites for all specificied RBPs on all targets and the lncRNA. Information derived from these predictions, as well as correlation data, is used to generate candidate mechanisms. Finally, the candidate with the lowest joint p-value is selected for each lncRNA-target pair, and a output list of mechanisms is produced. (*)Since at the time of publication only 22 RBP CLIP-Seq datasets were available for non-splicing related, post-transcriptional regulation proteins.

### 2.1 Sequence Decomposition by Accessibility

Since IntaRNA2 uses accessibility to predict RNA-RNA interactions, areas of low accessibility can be removed from the search space. An added benefit of this approach is that long transcripts can be naturally split into smaller sequences that can be analyzed independently. Since IntaRNA2 complexity increases quadratically with sequence length, sequence splitting makes cases tractable that are intractable otherwise, i.e., even transcripts with length *>*20kb can be considered. To accomplish a proper splitting, we developed a new algorithm that incrementally detects the least accessible (most structured) positions in the sequence to be used as split positions. The minimal number of splits are selected that are necessary to make every subsequence shorter than a user-specified length and for each of these subsequences to contain no position less accessible than its split positions. A default maximum length threshold of 1500 nt was selected to ensure that the memory usage does not exceed the typical amount of RAM on a PC or the per-core resource availability of a computing cluster. It should be noted that the majority of transcripts are less than the default threshold and therefore the heuristic will usually not be used, i.e., it is mainly applicable to extreme cases.

The algorithm finds the minimal set of most structured points at which to split a long input sequence according to a given length restriction as follows: Given a sequence *S*, the algorithm begins with position *x* = 0 and *y* = *|S| - l* where *l* is a fixed window length (IntaRNA seed length by default). First, the algorithm computes max_*i*_(*ED*(*i, i* + *l*)), where *x ≤ i ≤ y* and *ED* is the accessibility energy for that range. Accessibility energy is the energy required for a region of RNA to be single stranded, inversely proportional to the probability of the bases being paired in that region, and computed via the partition function. With the detected position *i*, a new interval (*i*+*l, |S|-*1) is created and put on the stack. Furthermore, for the current interval, *y* is updated to *i -* 1. This process is repeated until *y - x* + 1 is less than the length threshold, at which time it is added to the final list of intervals. The algorithm then moves to the next interval from the stack, i.e., the interval created in the last iteration. The iteration continues until the last interval is reached (the first interval created with endpoint *|S| - l*). Highly structured regions will produce many maximum *ED* windows in close proximity, so a minimum interval length is enforced (again, IntaRNA seed length by default) and regions shorter than this minimum are discarded. The final output is a set of intervals, which are then used as input for IntaRNA. More specifically, IntaRNA will sequentially go through each interval and find the optimal hybridization of the lncRNA with the subsequence contained within the interval. An example execution is shown in Figure 2.

**Fig. 2.**
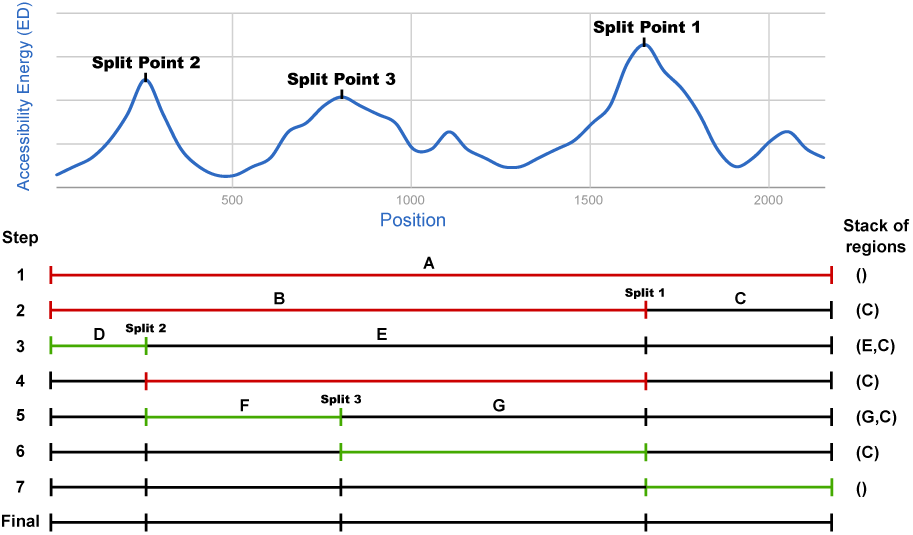
Example execution of the splitting algorithm with a max sequence length of 1000 nt, where the red interval is the one being processed. (1) The first iteration starts with the entire sequence which is longer than the threshold. (2) The first split occurs at the position with max ED at *∼* 1700 nt. (3) The interval is still too long, so a second split is made at the next position of max ED at *∼* 250 nt. (4) The interval is now below the threshold so the iteration continues to the next interval. (5) This interval is over the threshold and is split at *∼* 800 nt. (6-7) The next two intervals are below the threshold. (final) The end result is four intervals, all below the length threshold and more accessible than their split positions.

In a test with 100 random sequences of length over 1500 nt, the algorithm reduced the runtime by 13% and peak memory usage by 65%. Since peak memory has a constant upper bound when using this approach, the peak usage reduction is even more dramatic for extreme cases. It must be noted that the full accessibility matrix for the entire target/lncRNA structure is used for computing hybridization energies, and is reused for each interval within a target. This allows us to limit the search space of possible hybrids as described without any loss in optimality. In other words, any interaction calculated in an accessible region using subsequences of the input RNAs will be identical to those computed using the full input sequences. In the test above, out of the top 10% of predictions using the vanilla algorithm, 95% of them were identical to those found when using the decomposition. This number increases to 97% when we allow for small differences in predicted sites. The only case where an “optimal” interaction may be missed is if a highly energetic hybrid exists between highly structured regions of both RNAs where the difference in energy is still greater than the difference in energy for interactions in more accessible regions. It is unclear whether this type of interaction actually occurs in nature as such interactions exhibit slow kinetics.

### 2.2 RNA-RNA Interaction Predictions

The next stage is the prediction of RNA-RNA interactions using IntaRNA2 (Mann *et al*., 2017) with the modifications outlined above. Details on the IntaRNA2 algorithm can be found in Supplementary Section 1. IntaRNA2 is executed with the parameters `–tAccL 150 –tAccW 200 –qAccL 150 –qAccW 200 -n 5 –tRegionLenMax 1500`. The `AccL` and `AccW` options used by RNAplfold within IntaRNA2 are recommended by Lange et al. (Lange *et al*., 2012). The n option specifies the number of predictions (optimal + suboptimals). The `tRegionLenMax` option specifies the maximum length of an accessible sequence. This value was selected based on the available computational resources and the average RNA length in the reference transcriptomes. This reduces the usage of the heuristic to minimize the effect on the sensitivity of the algorithm.

MechRNA can run IntaRNA2 on a standard machine or distribute the computation across multiple jobs on a computing cluster. Interactions are predicted between the lncRNA and one of the two reference transcriptomes (Ensembl GRCh37.75 and GRCh38.86). The transcriptomes include all mRNA and ncRNA transcripts, excluding sequences less than 40 nt. This threshold was selected in order to include primary miRNA transcripts while removing dubious, unclassified transcripts. A subset of these transcriptomes is used if the user specifies a list of targets. Once all predictions are completed, the top most energetic interactions (default 3%) are selected for further analysis. P-values are computed for each of these interactions using a distribution estimated from the free energies of all interactions (details in Supplementary Section 2.1).

### 2.3 RNA-Protein Interaction Predictions

For determining RBP binding sites on transcripts, we rely on publicly available CLIP-Seq data. However, since CLIP-Seq depends on transcript expression, binding sites on transcripts specific to certain cell types or conditions cannot be recovered. As we want to study interactions across a reference transcriptome including lncRNAs specifically expressed in certain cancers, we would consequently miss many sites by relying only on direct binding evidence from CLIP-Seq. Therefore, to comprehensively capture protein binding information into our interaction models, we used GraphProt to create transcriptome-wide binding site predictions for 22 RBPs which are known to participate in post-transcriptonal gene regulation and influence transcript stability. As an example, using this approach we successfully predicted the interaction between hnRNP-L and the lncRNA DSCAM-AS1, for which there were no reads present in the hnRNP-L CLIP-Seq data (Niknafs *et al*., 2016). Based on the binding sites inferred from CLIP-Seq data for a given RBP, GraphProt learns its binding preferences and integrates these into a predictive model, incorporating either sequence (referred to as sequence model) or sequence and structure information combined (referred to as structure model). A detailed description of the algorithm can be found in Maticzka et al. (Maticzka *et al*., 2014).

For the 22 RBPs we trained 20 sequence and 8 structure models based on various CLIP-Seq data sources (Table 1). Models for each RBP were selected based on their performance in 10-fold-cross validation, preferring models with higher ROC (area under the receiver operating characteristic) and APR (mean average precision) values. The trained models were then used to predict nucleotide-wise binding score profiles (GraphProt setting: `-action predict_profile`) on two different reference transcriptomes (described in the previous section). Nucleotide-wise profile scores were further averaged with a sliding window approach, taking all scores up to 5 nt up-and downstream of the score position to calculate the new average score. Peaks were extracted from the average score profiles, where a peak is defined as the maximum score in a contiguous region of positive scores. In order to estimate score significancies and to make scores comparable between models, p-values for each peak score were calculated (details in Supplementary Section 2.2).

**Table 1.**
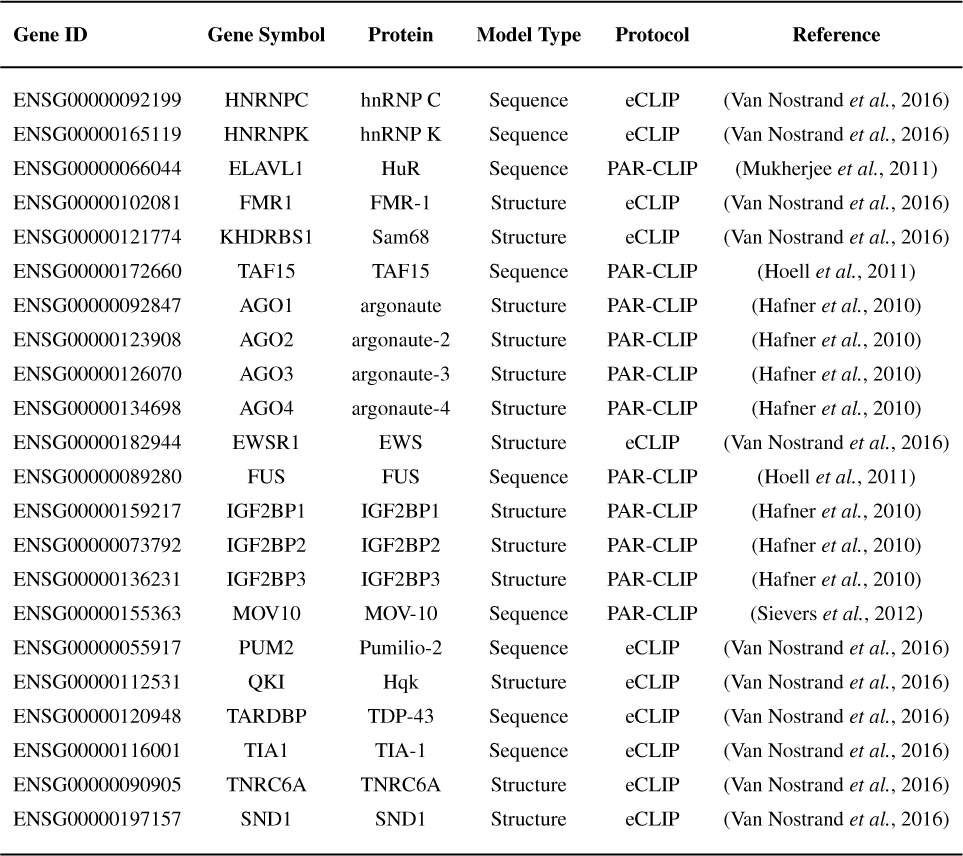
List of RBPs used in the analysis including the source CLIP-Seq data and model type.

### 2.4 Correlation Data from TCGA Protstate Tumor Samples

If screening mode is selected, correlation data is also incorporated for all RNA-RNA and RNA-protein pairs predicted in the previous stages. To obtain correlation data we used the GeneNet R package (Schafer and Strimmer, 2005). This approach first computes partial correlations for every pair of genes. The partial correlation is the correlation when the effects of all other variables (genes) are negated. These partial correlations are then used to create a graph where each edge is assigned a p-value. We used default parameters and a FDR cutoff of 0.2 to obtain the final correlation network. We deliberately allow a false discovery rate of 20% since the main information will be provided by the RNA-RNA and RNA-protein interactions.

The gene expression data used for correlation computation was derived from The Cancer Genome Atlas (TCGA) (Weinstein *et al*., 2013) patient samples. Specifically, this includes 551 RNA-Seq samples, 499 tumor and 52 normal. Only the tumor samples were used in the analysis. The raw read counts were normalized using DeSeq2 (Love *et al*., 2014). All genes with an average read count less than 1 were removed, resulting in 32,709 genes (coding/non-coding).

### 2.5 Combining Evidence

At this stage we incorporate the RNA-RNA and RNA-protein predictions in order to infer a potential mechanism for the lncRNA. For each target transcript, all combinations of RNA-RNA and RNA-protein interactions are classified into candidate mechanisms as shown in Figure 3. The number of combinations is reduced by considering the *a priori* information provided by the user and known functions of the RBPs (for example, HuR is primarily known to stabilize its bound RNA (Srikantan and Gorospe, 2012)). In screening mode, the correlations are also used at this stage to determine if a candidate mechanism is valid. For example, let a target RNA have a peak for RBP *A* and *B*, a lncRNA has a peak for *C*, and the RNA-RNA interaction between the two overlaps at the *A* peak. *A* is positively correlated with the target, *B, C*, and the lncRNA are negatively correlated with the target. Then the following tuples would be generated, where ([*target*_*peak*], [*lncRNA*_*peak*], [*mechanism*_*type*]) and a dash indicates absence of binding:

**Fig. 3.**
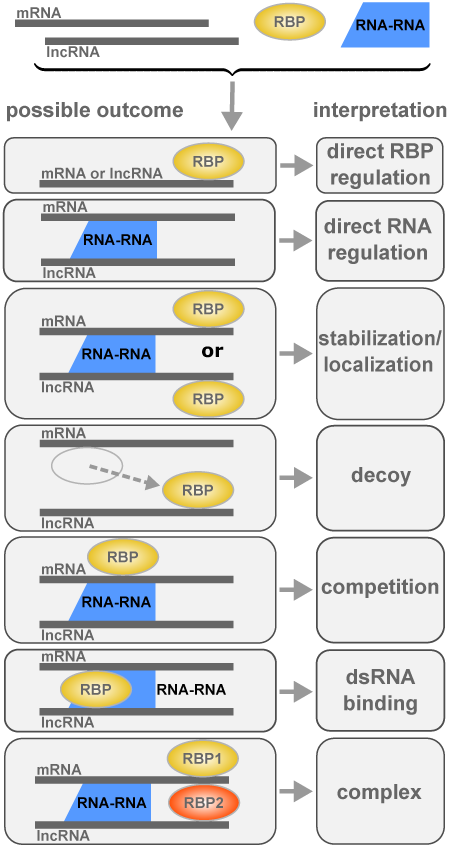
Illustration of the possible mechanisms that can be inferred from RNA-RNA and RNA-protein interactions.

- (*-, -, direct*_*downregulation*)
- (*A, -, competitive*_*downregulation*)
- (*-, C, localization*_*downregulation*)
- (*B, -, destabilization*)
- (*A, C, competitive*_*downregulation*)
- (*B, C, complex*_*downregulation*)

An explanation of each mechanism type with known examples is shown in Table 2. Decoy and direct RBP mechanisms are not included in the predictions since they do not include RNA-RNA interactions, making target prediction too non-specific (fully dependant on correlations). Double-stranded RNA binding mechanisms are not predicted either since the CLIP-Seq protocol does not capture such interactions.

**Table 2.** Descriptions of known lncRNA mechanisms. Mechanisms in italics are not included in the predictions.

| Mechanism | Description | Example |
| --- | --- | --- |
| **Direct RBP** | RBP interaction directly impacts the target or lncRNA | hnRNPL binding to DSCAM-AS1 (Niknafs *et al.*, 2016) |
| **Direct RNA** | RNA-RNA interaction directly impacts the target with no RBP involvement | TINCR stabilization of various mRNAs (Kretz *et al.*, 2013) |
| **(De-)Stabilization** | RNA-RNA interaction increases/decreases the affinity of RBP binding nearby | iNOS stabilization by AS via HuR (Matsui *et al.*, 2008) |
| **Localization** | RBP bound to the lncRNA is brought into the vicinity of the target through RNA-RNA interaction | MALAT1 localization of splicing factors (Bernard *et al.*, 2010) |
| **Decoy** | RBP is sequestered from the target by the lncRNA | Gas5-AS binding transcription factors (Kino *et al.*, 2010) |
| **Competitive** | RBP and lncRNA compete for the same binding location on the target | 7SL disrupts HuR stabilization of TP53 (Abdelmohsen *et al.*, 2014) |
| **dsRNA Binding** | A dsRNA binding protein interacts with stems created from lncRNA interaction | STAU1-mediated decay (Kim *et al.*, 2007) |
| **Complex** | The lncRNA facilitates the formation of a complex between multiple proteins | HOTAIR and the polycomb complex (Zhang *et al.*, 2014) |

The free energies of the RNA-RNA interactions and the peak scores of the RNA-protein interactions both have associated p-values. As mentioned before, each lncRNA-target and protein-target pair of correlations also has a p-value. These p-values can be used to quantitatively assess whether one mechanism is more likely than another. This requires the combining of up to six p-values (depending on the number of interactions involved) into a single p-value for each candidate. The intuitive way to accomplish this is to multiply the p-values together, however this is not correct since the product of p-values is not uniform under the null model. To solve this problem, we use the Stouffer’s Z-score method (Stouffer, 1949), which involves computing the sum of the inverse of a normal distribution of each p-value, followed by normalization. This approach also allows for weighting p-values, but we set all weights to be equal. The final output of the pipeline is the list of potential mechanisms sorted and filtered by the joint p-values.

## 3 Results and Discussion

We selected 8 lncRNAs to analyze using MechRNA, as summarized in Table 3. 7SL (Abdelmohsen *et al*., 2014), PCAT1 (Prensner *et al*., 2011) and ARlnc1 (Accepted in principle, Zhang et al. Nature Genetics 2018) recently investigated lncRNAs with known roles in prostate cancer and mechanistic hypotheses, are used to test the hypothesis-driven mode. The remaining 5 lncRNA are used to test the screening mode.

**Table 3.** Selected LncRNAs for MechRNA analysis. The lncRNAs vary in terms of what is known about their mechanisms, allowing MechRNA to be tested with various amounts of a priori data. PCAT1 has a question mark indicating that competitive binding is the hypothesis not been validated yet.

| lncRNA | Length | Target | Protein Binding | Mechanism | Cancer Type |
| --- | --- | --- | --- | --- | --- |
| 7SL | 299 | TP53 | HuR | Competitive | Prostate |
| PCAT1 | 1992 | BRCA2 | HuR | Competitive? | Prostate |
| ARlnc1 | 2786 | AR | Unknown | Unknown | Prostate |
| PCA3 | 3922 | Unknown | Unknown | Unknown | Prostate |
| PCAT29 | 694 | Unknown | Unknown | Unknown | Prostate |
| LINC00514 | 3385 | CLDN9 | Unknown | Unknown | NEPC |
| SSTR5-AS1 | 2864 | SSTR5 | Unknown | Unknown | NEPC |
| TINCR | 3733 | STAU1 | Many | Stabilization | Various |

PCA3 (Bussemakers *et al*., 1999) and PCAT29 (Malik *et al*., 2014) are well-studied prostate cancer related lncRNAs without a known mechanism. SSTR5-AS1 is one of the highest expressed lncRNAs in neuroendocrine prostate cancer (NEPC) and LINC00514 is one of the highest persistently expressed lncRNAs identified in the neuroendocrine transdifferentiation (NEtD) process, which is shown to cause NEPC (Ramnarine et al., unpublished). Finally we selected TINCR (Kretz *et al*., 2013) as a well known regulator of cell differentiation mediated by interaction with target mRNAs.

### 3.1 Hypothesis-driven Mode Results on Prostate Cancer lncRNAs

We first tested our hypothesis-driven mode with three prostate cancer lncRNAs. The first lncRNA is 7SL, which we use as a validation case since it has a good deal of evidence supporting the proposed mechanism. The next two lncRNAs, PCAT1 and ARlnc1, are less understood and so we aim to build a more complete picture of their potential mechanisms.

#### 3.1.1 7SL Downregulation of TP53 through Competitive Binding with HuR (ELAVL1)

Abdelmohsen *et al*. (Abdelmohsen *et al*., 2014) provided the first experimental evidence supporting a competitive lncRNA mechanism. 7SL is a housekeeping ncRNA which is part of the signal recognition particle (SRP) ribonucleoprotein complex, but also leads to increased cell proliferation when over-expressed in cancer cells. It was demonstrated that 7SL binds to the transcript of the tumor suppressor TP53 near HuR binding sites, preventing HuR from binding and subsequently reducing the stability of TP53. The experimentally validated RNA-RNA interaction was between nucleotide positions 10-56, 256-298 of 7SL and positions 2167-2300 of TP53 (ENST00000269305). Using PAR-CLIP data they determined that HuR binds at positions 2125-2160, 2452-2472 and 2531-2556.

For this case we ran MechRNA with 16 protein-coding TP53 transcripts as targets, all downregulatory mechanisms and all RBP models. For all 16 transcripts, “competitive downregulation” with HuR was predicted to be the most likely mechanism (*p <* 10^*-*15^ for the combined p-value as described in section 2.5). The predicted binding locations of 7SL and HuR for each transcript are shown in SupplementaryTable 2. The IntaRNA2 interaction prediction was in agreement with the crude BLAST search done in the experimental study. The 10-56 (actually 10-96 is more energetically favorable) interaction was also predicted but not included in the final results since it is not close enough to the HuR binding site to have an effect. In terms of RBP binding, GraphProt only predicts the 2125-2160 as significant when compared to all HuR binding across the transcriptome. This demonstrates the superiority of using GraphProt over raw PAR-CLIP data. We also show here that this mechanism appears to be ubiquitous across splice variants of TP53.

Another RBP, EWS, was included in this prediction. GraphProt detected a binding site for EWS on 7SL at 140-161, in between the two RNA-RNA interaction sites. EWS is best known for its role in Ewing sarcoma through its translocation with other genes. However, wildtype EWS also acts as a translation repressor by causing mRNA to be retained in the nucleus (Huang *et al*., 2014). It may be that EWS is aiding in the displacement of HuR and furthering the downregulation of TP53.

#### 3.1.2 PCAT1 Downregulation of BRCA2 through Competitive Binding with HuR (ELAVL1)

PCAT1 was identified by Prensner *et al*. (Prensner *et al*., 2011) as the most differentially expressed lncRNA in prostate cancer. Shortly afterwards it was discovered that this lncRNA regulates the important tumor suppressor BRCA2 (Prensner *et al*., 2014). Specifically, it was shown that PCAT1 reduces BRCA2 mRNA stability and that the first 250 nt of PCAT1 were essential for this process. Furthermore, they demonstrated that this regulation was occurring via the BRCA2 3’UTR. Since mRNA stability was decreased, our hypothesis is that a similar mechanism to 7SL exists for PCAT1 and BRCA2, so we used the same parameters of all downregulatory mechanisms and all RBP models. For this analysis we used the BRCA2 3’ UTR from the RefSeq transcript as it was used in Prensner *et al*. (Prensner *et al*., 2011).

Figure 4 summarizes the interaction predictions by showing the frequency of interaction for each position of PCAT1 and the significant RBP binding peaks. Our findings appear to support that the first 250 nt play an important role due to high frequency of interaction with targets an no significant binding with RBPs. The predicted mechanism was “competitive downregulation” (combined p-value *p <* 10^*-*4^) involving HuR on the 3’ UTR. The RNA-RNA interaction is between 11204-11237 on BRCA2 and 65-90 on PCAT1 (-12.493 kcal/mol), with a HuR peak at 11216-11236 on BRCA2. There are also two other HuR binding sites predicted by GraphProt downstream and upstream of the interaction site with similar binding affinity.

**Fig. 4.**
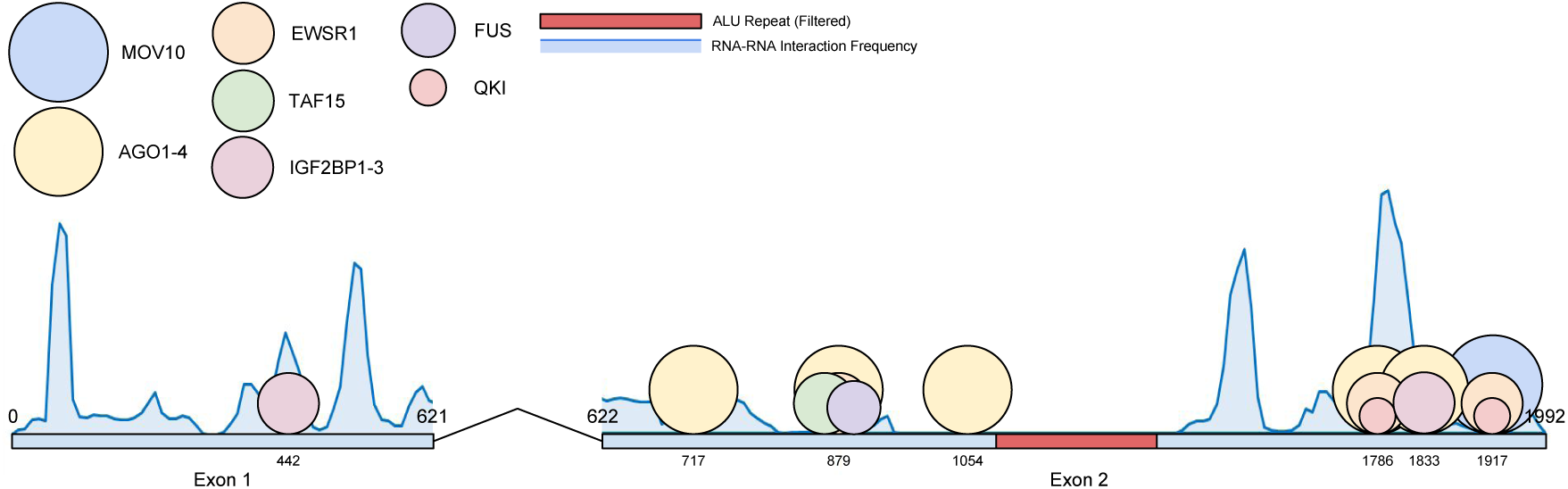
Distribution of RNA-RNA and RNA-protein interactions for PCAT1. The circles represent proteins and have a diameter relative to their molecular masses. The blue regions are a histogram of the frequency of RNA-RNA binding of each position with other transcripts.

To validate this mechanism experimentally, we first confirmed that HuR indeed binds to BRCA2 3’ UTR. As shown in Figure 5A, immunoprecipitation of HuR in LNCaP cells pulled down more BRCA2 mRNA than the IgG control. Next, we conducted a competitive binding assay in RWPE cells. This assay immunoprecipitated HuR using an anti-HuR antibody and the bound RNA (BRCA2) was detected by qPCR. In the presence of unmodified PCAT1 the amount of bound BRCA2 RNA was reduced. When using a modified PCAT1 construct with the first 250 nt deleted, there was no affect on the amount of bound BRCA2. This suggests that an interaction involving the 5’ end of PCAT1 is competitively reducing the amount of HuR bound to BRCA2 (Figure 5B).

**Fig. 5.**
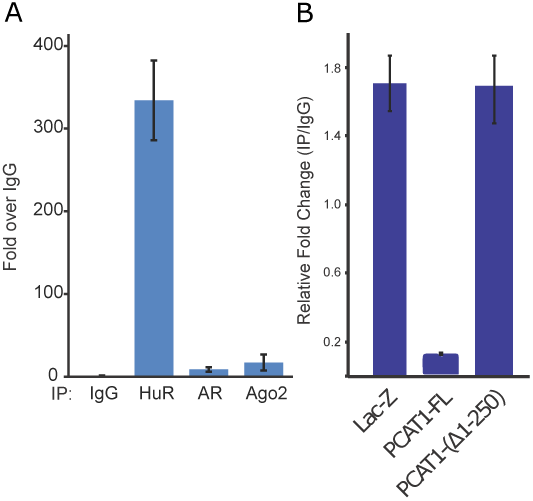
(A)IgG/HuR/AR/Ago2 proteins were immunoprecipitated by antibodies in LNCaP cells and the bound BRCA2 RNA was detected by qPCR. The result confirms binding of BRCA2 mRNA to HuR protein in cells. (B) RWPE cells stably expressing lac-Z, PCAT1-FL or PCAT1-delta-1-250 were harvested. HuR was immunoprecipitated using anti-HuR antibody and bound RNA (BRCA2) was detected by qPCR. As shown in A, HuR can bind to BRCA2. In presence of FL-PCAT1 this binding is inhibited. In presence of PCAT1-delta-1-250 there was no effect on HuR and BRCA2 binding. The result confirms the role of PCAT1 in mediating BRCA2-HuR binding.

#### 3.1.3 ARlnc1 Upregulatory Feedback Loop with Androgen Receptor

ARlnc1 has recently been identified as an upregulator of Androgen Receptor (AR) in prostate cancer (Accepted in principle, Zhang et al. Nature Genetics 2017). In turn, AR upregulates ARlnc1, leading to a positive feedback loop contributing to cancer progression. The mechanism was identified with the aid of the first stage of MechRNA, which predicted an RNA-RNA interaction between ARlnc1 and the 3’UTR of AR. However, how exactly ARlnc1 upregulates AR remains unclear. Similarly to 7SL, we ran MechRNA with all RBPs on all AR protein coding transcripts, but with all upregulation mechanisms.

The most common and important AR transcript, ENST00000374690, as well as two other splice variants (ENST00000612452, ENST00000396044) had predicted mechanisms involving the experimentally validated interaction at 815-851 on ARlnc1 (-35.8 kcal/mol). In all three cases, a “stabilization” mechanism was predicted (respectively *p <* 10^*-*6^, *p <* 10^*-*5^, *p <* 10^*-*4^ for the combined p-values) involving the protein Sam68, which has a strong binding site upstream of the ARlnc1 interaction on the AR 3’UTR. In agreement, Sam68 3’UTR interaction has been shown to enhance target translation (Paronetto *et al*., 2009). Sam68 is known to increase AR-V7 (ENST00000504326) expression (Stockley *et al*., 2015), but the authors observed that upregulation of AR-V7 (and full-length) was still present when using a mutated exonic splicing enhancer (ESE) site. They suggested a synergistic stabilization mechanism via the 3’UTR. Although the 3’UTR of AR-V7 and full-length AR is not shared, a similar binding pattern is observed for Sam68 and ARLnc1 in the AR-V7 3’UTR. Our findings appear to support the additional stabilization mechanism they observed and that all major AR isoforms are regulated in the same manner.

### 3.2 Screening Mode Results on Prostate Cancer lncRNAs

We ran MechRNA on all eight lncRNAs (three from the hypothesis-driven analysis and five additional cancer-related lncRNA as described in Table 3) using the entire transcriptome for potential targets for a broad, unbiased screen. This yielded several hundred to several thousand potential targets for each lncRNA. The number of predictions increased with the length of the lncRNA, since longer lncRNAs are more likely to have RNA-RNA and RNA-protein interactions and consequently more viable combinations of interactions, indicating potential mechanisms. Since our focus here is on cancer, we extracted predicted mechanisms involving known cancer genes from the TSGene (Zhao *et al*., 2015) and ONGene (Liu *et al*., 2017) database. These mechanisms are shown in Table 4.

**Table 4.**
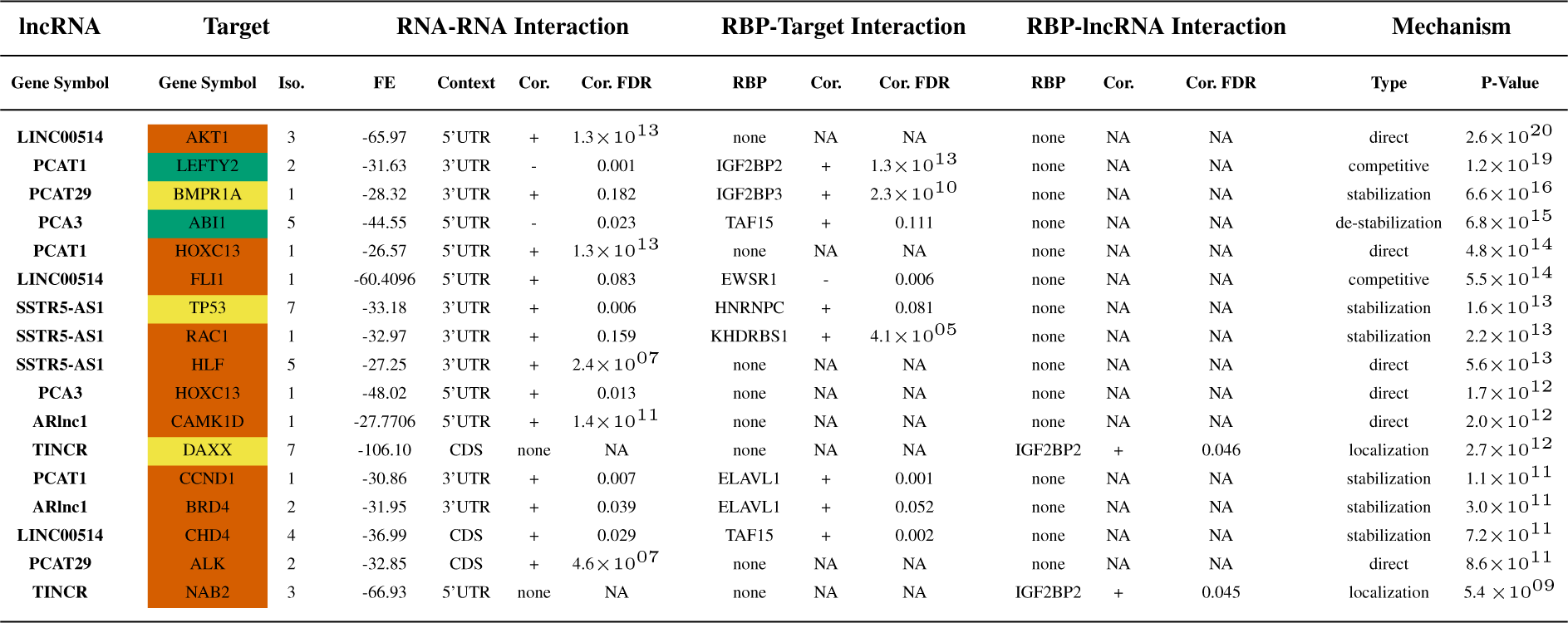
Select lncRNA mechanisms predictions for known cancer genes, selected based on rank (joint p-value) and agreement with known roles of the cancer genes and RBPs. Genes in red indicate oncogenes, green indicate tumor suppressors and yellow are uncategorized. The first section indicates the target and how many isoforms (Iso) it interacts with. The next three sections describe the interactions involved. For RNA-RNA, the free energy in kcal/mol (FE) and genomic context are included. For RBP-RNA, the protein name is provided. In all three cases the correlation (+ positive, -negative) and the correlation FDR are shown if applicable. The final section displays the mechanism categorization and the combined p-value.

As shown in the table, these prostate cancer lncRNAs generally act as positive regulators of oncogenes with the exception of the PCA3-ABI1 and PCAT1-LEFTY2 interaction. Also the most favorable RNA-RNA interactions commonly occur in the 5’ and 3’ UTRs, as would be expected for post-transcriptional regulation. It is unclear whether the CDS interactions have any functionality. TINCR-DAXX falls within a small simple repeat region which may indicate non-specific binding. Another observation is that PCAT1 and PCA3 share the target gene HOXC13, and even bind to the same location on the HOXC13 transcript. HOXC13 is commonly dysregulated in prostate cancer (Komisarof *et al*., 2017). It may be that the same phenotype is induced by both lncRNAs.

Our most significant result is an interaction involving AKT1, an important and well-studied prostate cancer gene (Cariaga-Martinez *et al*., 2013). LINC00514 binds very strongly to the 5’UTR and has a strong positive correlation, implying a direct upregulatory effect. No significant protein binding was detected in the region for the included proteins. This would suggest the lncRNA alone is able to regulate AKT1. We observed several other cases like this, labelled as “direct” in the table. It may be the case that some other RBP, which was not included in our analysis due to missing CLIP data, also interacts with AKT1 in this region. As the number of RBPs with available CLIP data is ever increasing, it is likely that a future run of MechRNA with more RBPs might provide additional evidence.

Another significant result was the predicted competitive downregulation of LEFTY2 by PCAT1. This is the most significant result for PCAT1 involving a known cancer gene. It has a close similarity to the PCAT1-BRCA2 mechanism, as it involves the same part of PCAT1 (61-97) binding to a 3’UTR overlapping a protein binding site (in this case IGF2BP2). LEFTY2 is an important tumor suppressor in endometrial cancer (Alowayed *et al*., 2016). We do not have data for PCAT1 expression in endometrial cancer, but there is high expression in ovarian and breast cancer (Iyer *et al*., 2015).

The PCA3-ABI1 mechanism is an interesting example demonstrating the importance of sequence accessibility for interaction prediction. ABI1 is known to negatively regulate cell growth and transformation and is down-regulated in a variety of cancers (Zhang *et al*., 2015; Chen *et al*., 2010; Cui *et al*., 2010). The gene has 11 annotated protein-coding isoforms in Ensembl, 9 of which have an identical 5’UTR sequence. However, 3 of the splice variants exclude exon 3, leading to a much more energetic binding to PCA3 (-13 kcal/mol difference). This is because the exclusion affects the accessibility of the 5’UTR by reducing the probability that this region is bound by intramolecular interactions. If PCA3 is indeed down-regulating ABI1, as the correlation indicates, there may be selection for these isoforms in cancer cells to increase the effect of PCA3. Naive approaches to RNA-RNA interaction prediction computing only the hybridization would not capture the difference in interaction energy between the different splice variants. This is because the sequence of the best hybridization site is always the same, the only feature considered when computing the optimal interaction. However, the accessibility can differ between different isoforms, which may affect the location of the true optimal interaction site, as we see in the case of PCA3-ABI1.

## 4 Conclusion

Recent discoveries of lncRNA mechanisms indicate that there exists a complex interplay between RNA-binding proteins, lncRNAs and their target RNAs. Until now, RNA-RNA and RNA-protein interaction predictions were carried out independently, failing to capture this complexity. Here we present MechRNA, the first tool to integrate both kinds of interactions in order to more accurately predict lncRNA mechanisms. We accomplish this by combining the output of IntaRNA2 and GraphProt into a novel inference tool which determines the most likely combination of interactions. These sets of interactions are then classified into mechanisms using correlation data from publicly available patient gene expression samples or user-defined *a priori* data. We demonstrated the functionality of MechRNA by analyzing 8 prostate cancer lncRNAs with varying amounts of information available with respect to their mechanisms. The results confirm one known mechanism, provide new insights into poorly understood mechanisms and offer new hypotheses for the remaining lncRNAs without known mechanisms. Despite the challenges involved in this kind of analysis (discussed in Supplementary Section 3), our results show that MechRNA is a useful tool for identifying potential roles of lncRNAs in cancer, and for furthering our understanding on lncRNA mechanisms in general.

